# Epigenetic variation in the Lombardy poplar along climatic gradients is independent of genetic structure and persists across clonal reproduction

**DOI:** 10.1101/2022.11.17.516862

**Authors:** Bárbara Díez Rodríguez, Dario Galanti, Adam Nunn, Cristian Peña-Ponton, Paloma Pérez-Bello, Iris Sammarco, Katharina Jandrasits, Claude Becker, Emanuele De Paoli, Koen J.F Verhoeven, Lars Opgenoorth, Katrin Heer

## Abstract

- Environmental changes can trigger phenotypic variation in plants through epigenetic mechanisms, but strong genetic influences make it difficult to isolate and study epigenetic effects. Clonal trees with low genetic variation, such as the Lombardy poplar *(Populus nigra* cv. ‘Italica’ Duroi), offer a unique system to study epigenetic variation associated with the environment.
- We collected cuttings (ramets) of Lombardy poplar along a wide geographical range in Europe. We performed whole-genome-bisulfite sequencing of 164 ramets grown in a common garden and of a subset of 35 of the original parental individuals. Using historical bioclimatic data, we tested the relationship between DNA methylation and climatic gradients.
- We found that average methylation levels in TEs and promoter regions correlate with biologically relevant climatic variables. Furthermore, we observed that DNA methylation was transmitted to the next clonal generation, but a fraction of the methylome changed relatively fast when comparing the parental individuals with the clonal offspring.
- Our results suggest that the poplar methylome is a dynamic layer of information that can be transmitted to the clonal offspring and potentially affect how poplars acclimate to new environmental conditions.

## Introduction

In the last couple of decades, extreme weather events have been increasing, often exceeding plants’ and animals’ tolerance thresholds, and driving mass mortalities in many species (IPCC, 2022). Understanding how plants respond to such weather events and other environmental conditions has thus become crucial for conservation policies and forest management programs. In studies on plant natural populations, intraspecific genetic diversity has been shown to contribute to the resistance and resilience of populations (Hughes et al., 2008). Genetic variation provides the baseline for phenotypic variation on which evolutionary processes can act, and plays an important role in plant adaptation (Fisher, 1958; Hughes *et al*., 2008). However, advances in molecular biology and genomics have shown that phenotypic variation among individuals is not only determined by genetic variation (Rapp & Wendel, 2005). One additional cause of phenotypic variation is epigenetic variation (Cubas *et al*., 1999; Manning *et al*., 2006; Xie *et al*., 2015). Several studies have shown that epigenetic variation can be spatially structured among and within plant populations, and that such a structure can be associated with environmental variation and phenotypic differentiation (Lira-Medeiros et al., 2010; Medrano et al., 2014; Avramidou et al., 2015; Kawakatsu et al., 2016; de Kort et al., 2020; Boquete et al., 2021; Galanti et al., 2022, Sammarco et al., 2022). Although causal relationships remain to be studied, such observations suggest that epigenetic variation could contribute to the acclimation of plants to changes in environmental conditions.

There are several molecular mechanisms involved in epigenetic variation, such as histone modifications, DNA methylation and small RNA-mediated processes (reviewed in Lloyd and Lister, 2022). Among these, DNA cytosine methylation (mC), is currently the most widely studied and best characterized modification (Zemach *et al*., 2013; Matzke & Mosher, 2014; Zhang et al., 2018; Lloyd & Lister, 2022) and consists of a base alteration in which a methyl group is added to the 5th carbon of a cytosine (Moore *et al*., 2012). In plants, cytosine methylation occurs at three different sequence contexts: CG, CHG and CHH, where H = A, T or C. Methylation at the CG and CHG contexts is usually symmetrical across both DNA strands, whereas methylation at CHH sites is asymmetrical (Meyer et al., 1994; Finnegan et al., 2003; Zhang et al., 2006; Lister et al., 2008). As a result of different mechanisms involved in DNA methylation maintenance, different sequence contexts differ in their degrees of *mitotic stability,* which are mainly dictated by their symmetry. In the symmetrical contexts, methylation maintenance is guided by the complementary DNA strand, and thus stably inherited across mitotic divisions (Niederhuth & Schmitz 2014). On the other hand, methylation in the asymmetrical context is maintained mainly by *de novo* establishment and thus less stable across cell divisions (Peter Meyer & Lohuis, 1994). In addition, depending on the genomic feature context, DNA methylation has different roles. For example, methylation in all sequence contexts is associated with silencing of transposable elements (TEs), while CG methylation is found in promoters of transcriptionally inactive genes and in the gene body of active genes (reviewed in Niederhuth & Schmitz, 2017). Variation in DNA methylation can be under genetic control (Zhang *et al*., 2018; Johannes & Schmitz, 2019) and arise stochastically as a result of imperfect DNA methylation maintenance (Becker *et al*., 2011; Schmitz *et al*., 2011; Johannes & Schmitz, 2019), or be induced by environmental conditions (Raj *et al*., 2011; Bräutigam *et al*., 2013; Lämke & Bäurle, 2017). Furthermore, some of these methylation marks can be transmitted from parental individuals to offspring (Johannes *et al*., 2009; Becker & Weigel, 2012; Herman & Sultan, 2016; Gáspár *et al*., 2019; Boquete *et al*., 2021). If DNA methylation can be induced by environmental conditions, we would expect patterns of DNA methylation to be associated with geographic or climatic gradients beyond what can be explained by the underlying genetic structure of the studied population. Several studies indeed found correlations between methylation patterns and habitat or climate in different plant species. However, almost all these studies were conducted on sexually reproducing plant species, were constrained to small-scale geographic gradients, or used low-resolution molecular methods (Lira-Medeiros *et al*., 2010; Nicotra *et al*., 2015; Avramidou *et al*., 2015; Gugger *et al*., 2016; Herrera *et al*., 2017; Gáspár *et al*., 2019). With the continuous decrease of sequencing costs, recent studies based on whole genome bisulfite sequencing (WGBS) have provided more detailed methylation data (Dubin *et al*., 2015; Kawakatsu *et al*., 2016; de Kort *et al*., 2020; Galanti *et al*., 2022). With WGBS we can now quantify methylation at the scale of whole genomes and accurately map methylated cytosines at a single-base resolution (Lister and Ecker, 2009). Nevertheless, the extent to which genetic variation influences epigenetic variation is still not clear (Richards *et al*., 2010, 2017). Studing epigenetic variation in asexually (i.e. clonally) reproducing species allows focusing on epigenetic variation in the absence of confounding genetic variation. Moreover, during sexual reproduction, some proportion of the methylation patterns might be reset (Wibowo et al., 2016), whereas we assume that they are faithfully transmitted during clonal propagation. Thus epigenetic marks have therefore the potential to be stably transmitted across clonal generations and may thus create heritable phenotypic variation (Verhoeven & Preite, 2014).

Since the first assembly of the *P. trichocarpa* genome in 2006, the amount of available genetic, genomic, and biochemical resources have increased considerably, and *Populus* species have become a model for studying plant adaptation (Taylor, 2002; Tuskan et al., 2006; Jansson & Douglas, 2007). The Lombardy poplar *(Populus nigra* cv. ‘Italica’ Duroi) is a widely distributed tree clone. This variety likely originated in the 18th century from one single male tree of *P. nigra,* located in central Asia (Elwes & Henry, 1913), and was spread by cuttings worldwide from Italy. It is assumed that most Lombardy poplars originate from artificial propagation performed by humans (CABI, 2022).

Here, we present the first study investigating DNA methylation variation in a clonal tree species. We collected poplar cuttings from a wide climatic and geographic gradient across Europe and planted them in a common garden in Central Germany. We analyzed methylation variation among trees in the field and in the common garden. Thus we were able to address two questions: (1) given a uniform genetic background, do different environmental conditions result in differences in DNA methylation in Lombardy poplar? If so, (2) do these differences persist over time after clonal propagation in a common environment?

## Materials and Methods

### Plant material and common garden design

Between February and March 2018, we sampled cuttings from *Populus nigra cv* ‘Italica’ clones in Europe across geographical gradients that spanned from 41° to 60° N and −5° to 25° E approximately, at twelve sampling sites that covered seven different Köppen-Geiger climate subtypes (Peel *et al*., 2007). We tagged and georeferenced the source trees (hereafter referred to as “ortets”). During the first week of May 2018, we planted the cuttings (hereafter referred to as “ramets”) in a common garden in the Marburg Botanical Garden (Germany) under a random block design. The common garden area was not shaded in any way, allowing the ramets to grow under direct sunlight. No herbicides, pesticides, or fertilizers were used in the common garden. We planted the ramets with 1 m between trees and watered them frequently for a period of five months until the end of summer. A more detailed description of sampling and the common garden set-up can be found in Díez Rodríguez et al., (2022).

### Whole genome bisulfite sequencing

Of the 375 individuals considered to belong to the same genotype by Díez Rodríguez *et al.* (2022), we selected a subset for WGBS. We chose 14 ramets from 12 sampling sites from the common garden, except for those from Lithuania, of which only 10 ramets had survived in the garden, resulting in a total of 164 individuals. From the original set of ortets, we chose 5 individuals from seven out of the 12 sampling sites, with a total of 35 individuals. We collected leaf material from individuals, both in the field and in the common garden, at approximately the same time in July 2018. We extracted DNA from leaf tissue obtained from mature, healthy leaves dried in silica gel using the PeqGOLD Plant DNA mini kit (PEQLAB Biotechnologie GmbH, Erlangen, Germany). We used the NEBNext Ultra II DNA Library Prep Kit for sequencing library preparation, combined with EZ-96 DNA Methylation-Gold MagPrep (ZYMO) for bisulfite libraries. The protocol involved: i) end repair and 3’ adenylation of sonicated DNA fragments, ii) NEBNext adaptor ligation and U excision, iii) size selection with AMPure XP Beads (Beckman Coulter, Brea, CA), iv) bisulfite treatment and cleanup of libraries, v) PCR enrichment and index ligation using Kapa HiFi Hot Start Uracil+ Ready Mix (Agilent) for bisulfite libraries (14 cycles), vi) final size selection and cleanup. Finally, we sequenced paired-end for 150 cycles on a HiSeq X Ten instrument (Illumina, San Diego, CA). All sequenced raw fastq files are available at the European Nucleotide Archive (ENA) database, under project number PRJEB44879.

### Methylation data and DMR calling

For the methylation analysis we used the EpiDiverse toolkit (version 1.0), a pipeline suite for WGBS data analysis in non-model plant species (Nunn *et al*., 2021). For alignment, quality control, and methylation extraction we used the WGBS pipeline. This pipeline uses FastQC (https://www.bioinformatics.babraham.ac.uk/projects/fastqc/) to perform quality control, erne-bs5 (Prezza *et al*., 2012; http://erne.sourceforge.net/) to map raw reads, Picard MarkDuplicates (https://broadinstitute.github.io/picard/) to filter PCR duplicates and MethylDackel (https://github.com/dpryan79/MethylDackel) to perform the methylation calling. We mapped the samples to the *Populus nigra* cv ‘Italica’ reference genome, freely available at the European Nucleotide Archive (ENA) under project number PRJEB44889. We only retained uniquely-mapping reads longer than 36 bp. On average, around 80% of the total number of reads were mapped to the reference genome. We calculated the bisulfite non-conversion rate using the mitochondrial genome, and found a mean rate of 0.005. Mapping stats and conversion rates for each individual sample are shown in Supplementary Table 1. Methylation levels for each called position were calculated according to Schultz et al. (2012) and using the following formula (*C* = reads supporting methylated cytosine, *T* = reads supporting unmethylated cytosine, *i* = position of cytosine):

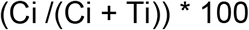

We obtained individual bedGraph files for each sample and context. We filtered out positions with a coverage lower than 6. For five ramet samples, less than 60% of the initial positions remained after filtering, and were thus excluded from the data set. We then merged the individual files into multisample bed files using custom scripts based on the *unionbedg* command from the BEDTools suite (Quinlan & Hall, 2010), retaining positions that were called in at least 80% of the samples. To directly compare only positions with methylation calls common to all samples, we obtained three different files per context. The first file contained 35 ortet samples (as mentioned in the plant material section); the second file contained 158 ramet samples; and lastly, the third file contained 64 paired ortet and ramet samples (32 samples from ortets and 32 from their respective ramets). A summary of the number of samples and the number of positions retained in each file is shown in Table 1. To study the epigenetic structure of the poplar clones, we ran Principal Component Analysis (PCA) per context using the *prcomp* function of the *stats* package (ver. 4.1.3; R Core Team, 2022).

**Table 1.** Summary of number of samples and positions included in each file used for methylation analysis

| Type of file | N Samples | N positions per context |  |  |
| --- | --- | --- | --- | --- |
|  |  | CpG | CHG | CHH |
| Ortet | 35 | 8,318,522 | 13,678,685 | 76,501,469 |
| Ramet | 158 | 7,820,008 | 12,961,553 | 72,754,297 |
| Paired | 64 | 8,139,896 | 13,412,560 | 75,215,708 |

The EpiDiverse toolkit (Nunn *et al*., 2021) includes a DMR pipeline that uses metilene (Jühling *et al*., 2016) to call Differentially Methylated Regions (DMRs) between all possible pre-defined pairwise comparisons between sites for each sequence context. We used the default parameters of the DMR pipeline to define DMRs. In this study, each sampling site where the ramets were collected was considered as an individual group and compared to all the other sites. DMRs were called among three different group sets. First, we ran the DMR pipeline using only groups containing ortet samples in each pairwise comparison; second, we compared groups containing only ramet samples; and third, we compared ortet samples with their paired ramet samples. We then used custom scripts to summarize the results of the pipeline, and obtained a single file for each context and each run with a list of all DMRs, their genomic coordinates, and the specific pairwise comparison they belonged to. Supplementary Figure 1 shows a schematic description of the pairwise comparison design.

### Variant calling, filtering and imputation

We used the EpiDiverse SNP pipeline (Nunn *et al*., 2021, 2022) with default parameters to infer Single Nucleotide Polymorphisms (SNPs) from WGBS data. We combined the output of individual Variant Call Format (VCF) files from the ramet samples into a multisample VCF file using BCFtools (*v1.9*, Danecek et al., 2011). We filtered for variants successfully genotyped in at least 90% of individuals, with a minimum quality score of 30 and a minimum mean depth of 3. For the PCA analysis, we retained only biallelic SNPs and removed SNPs with more than 10% missing values and a Minor Allele Frequency (MAF) < 0.01. The remaining missing values were imputed with BEAGLE v 5.1 (Browning, Zhou, and Browning 2018). We also removed SNPs that were heterozygous in more than 95% of the samples. To reduce the number of SNPs for downstream analysis, we filtered redundant SNPs by pruning for Linkage Disequilibrium (LD) with a maximum LD of 0.8 between SNP pairs in a sliding window of 50 SNPs. After filtering and imputing, we were able to retain 343,977 SNPs. We performed the PCA analysis with PLINK *(v1.90b6.12,* Purcell et al., 2007) and plotted the results with custom scripts in R (https://github.com/EpiDiverse/scripts).

### Correlation between methylation and bioclims

To assess correlations between methylome variation and climatic variables, we obtained bioclimatic data for each of the locations of the ortets from the CHELSA timeseries data set (Karger *et al*., 2017). The CHELSA data set covers the period between 1979 and 2013 and provides gridded data at a resolution of 30 arcsec (~ 1km). We included all 19 bioclimatic variables, as described in the CHELSA web page: https://chelsa-climate.org/bioclim/. Bioclimatic data for all sequenced individuals is available in Zenodo at https://doi.org/10.5281/zenodo.5995424. The methylation data for specific genomic regions used in the correlation analysis was obtained using the BEDTools *intersect* command (Quinlan and Hall, 2010) and a custom structural annotation. The annotations are available at the European Nucleotide Archive (ENA) under project number PRJEB44889. We correlated average global methylation levels with CHELSA bioclims using the Spearman method. The analysis was performed with the *corr.test* function of the *psych* package (ver. 2.2.5, Revelle, 2022) and plotted using the *heatmap.2* function of the *gplots* package (ver. 3.1.3, Warnes et al., 2022).

### Mantel tests

To investigate if epigenetic distance between individual ramets was correlated with geographic, climatic and/or genetic distance, we performed mantel tests, using the *mantel* function of the *vegan* package (ver. 2.5-7; Oksanen et al., 2013). As input for the geographic and climatic distance matrices, we used the original geographic coordinates and the bioclimatic data of the ramets. We calculated two types of epigenetic distance matrices. The first matrix was based on the methylation levels of single methylated positions (MPs). In the second matrix, we used the BEDTools suit to merge the DMRs called from multiple pairwise comparisons in order to obtain a union set of candidate regions, variable between two or more populations of ramets. We then calculated mean methylation levels (according to Schultz et al. 2012) in each region. For the genetic distance matrix we used the same SNPs that were used for the genetic structure analysis. To standardize the data and make it comparable, we then conducted a PCA and calculated the first three PCs for each type of input. We then created Euclidean distance matrices using the *dist* function of the R *stats* package (Version 4.2.1, R core team, 2022). Finally, we ran the mantel tests with the Pearson correlation method and 9999 permutations.

### Persistence of DNA methylation patterns

To study if methylation patterns were conserved across clonal generations, we focused on the seven sites for which we had collected samples from ortets and ramets. We called DMRs between sites for ortets, for ramets, and between ortets and ramets from each site. Supplementary Figure S2 shows the total number of DMRs for each pairwise comparison among ortets (A) and ramets (B), respectively, ordered according to latitude of origin from South to North. If methylation patterns are conserved in the next clonal generation we assumed we would be able to find the same DMRs when comparing the same sampling-site pairs between ortets and between ramets. We therefore intersected the bed files with all the DMRs called using the BEDTools intersect command. Specifically, we intersected a file containing DMRs called from group A vs group B ortets with a file containing DMRs called from ramets belonging to the same groups (i.e. corresponding to the clonal offspring). We then repeated the analysis for each of the 21 possible pairwise comparisons between sites. Supplementary Figure S4 shows a detailed count of hypermethylated and hypomethylated DMRs for each pairwise comparison. After running the intersections, we created individual files containing all the regions found among ramets that overlap with regions found among ortets.

## Results

### Methylation profiles in the Lombardy poplar

Average global methylation in ramets of the Lombardy poplar from 12 different sampling sites ranged from 30 to 40% in the CG context, 15-25% in the CHG context and 1-3% in the CHH context (Figure 1A). We did not find any statistically significant differences among methylation levels from different sites in any of the contexts, and variation within each group seemed to be higher than the variation among groups. We found the highest number of DMRs in the CHG context (~130,000 DMRs), followed by the CG context (~ 70,000 DMRs) and the CHH context, where only ~ 9,100 DMRs were called among all pairwise comparisons among sites (Figure 1B). However, most of these DMRs were common to two or more comparisons. When common DMRs were merged into unique regions, we found around 11,400 CG-DMRs, 14,400 CHG-DMRs and 4,100 CHH-DMRs. The length of the merged DMRs ranged from 10 to around 5,000 bases (Supplementary figure S3). Of these DMRs, a considerable fraction overlapped with annotated transposable elements (TEs) in all sequence contexts (~4,600; ~ 10,500 and 4,200, respectively for CG, CHG and CHH). Interestingly, only 31 DMRs in the CHH context overlapped with coding sequences (CDS), while around 4,600 CG-and 3,100 CHG-DMRs overlapped with these regions (Figure 2c).

**Figure 1.**
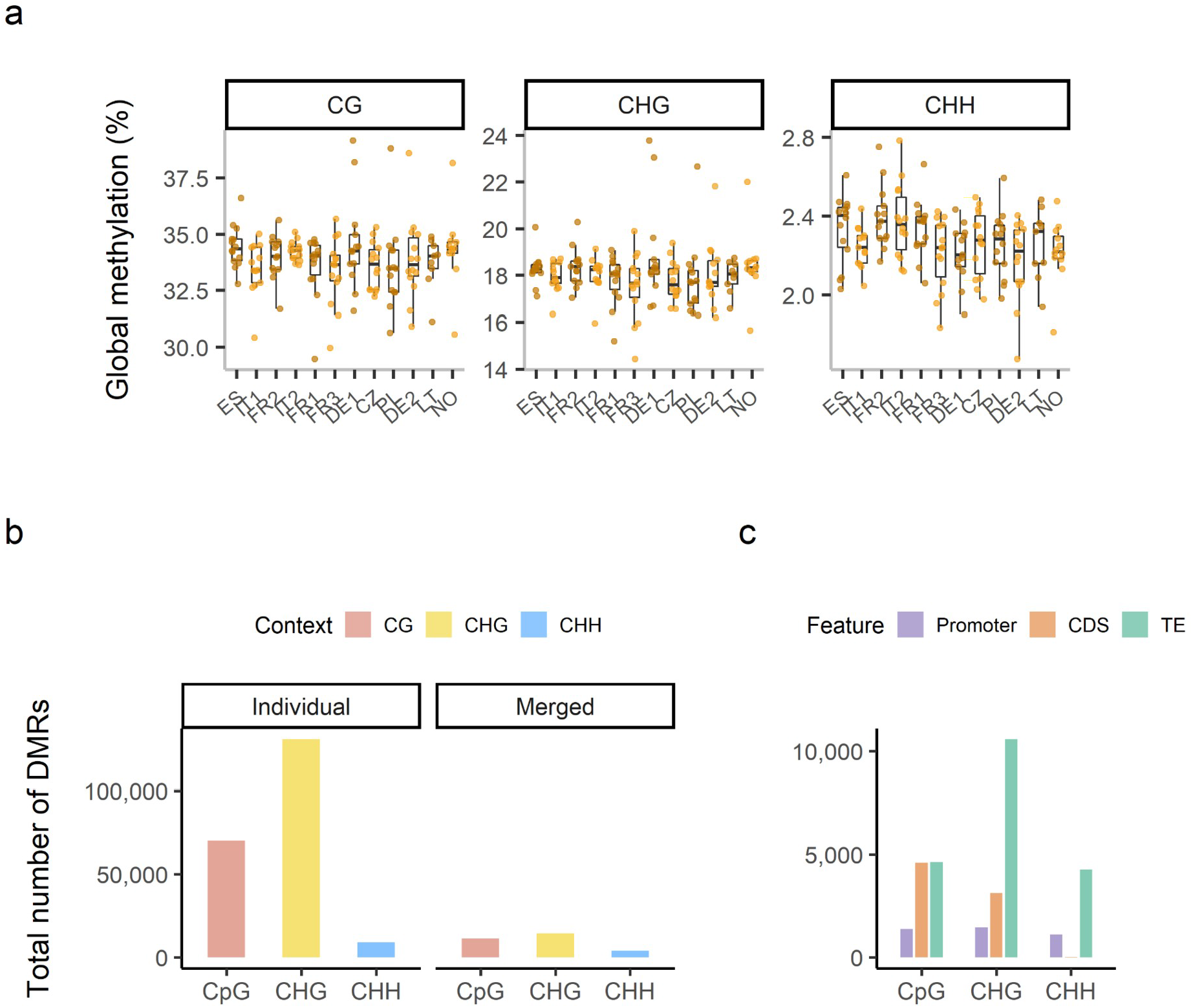
Methylation profiles in the Lombardy poplar (ramets). **a**. Variation in methylation levels among ramets across geographical gradients in all sequence contexts. Sites are ordered from South to North according to their geographic coordinates and labeled by the sample site code (ISO 3166 standard country code): ES: Spain, n = 14; IT1: Italy 1, n = 13; FR2: France 2, n = 13; IT2: Italy 2, n = 14; FR1: France 1, n = 14; FR3: France 3, n = 14; DE1: Germany 1, n = 13; CZ: Czech Republic, n = 14; PL: Poland, n = 14; DE2: Germany 2, n = 14; LT: Lithuania, n = 9; NO: Norway, n = 12. Note the different scales in the Y axes (n = 158). **b.** Total number of DMRs in each sequence context, called from all pairwise comparisons (n = 158). Number of individual DMRs is the sum of DMRs obtained from all pairwise comparisons. Total number of merged DMRs corresponds to regions that were merged into unique DMRs **c.** Total number of merged DMRs overlapping specific genomic features in each sequence context (n = 158).

**Figure 2.**
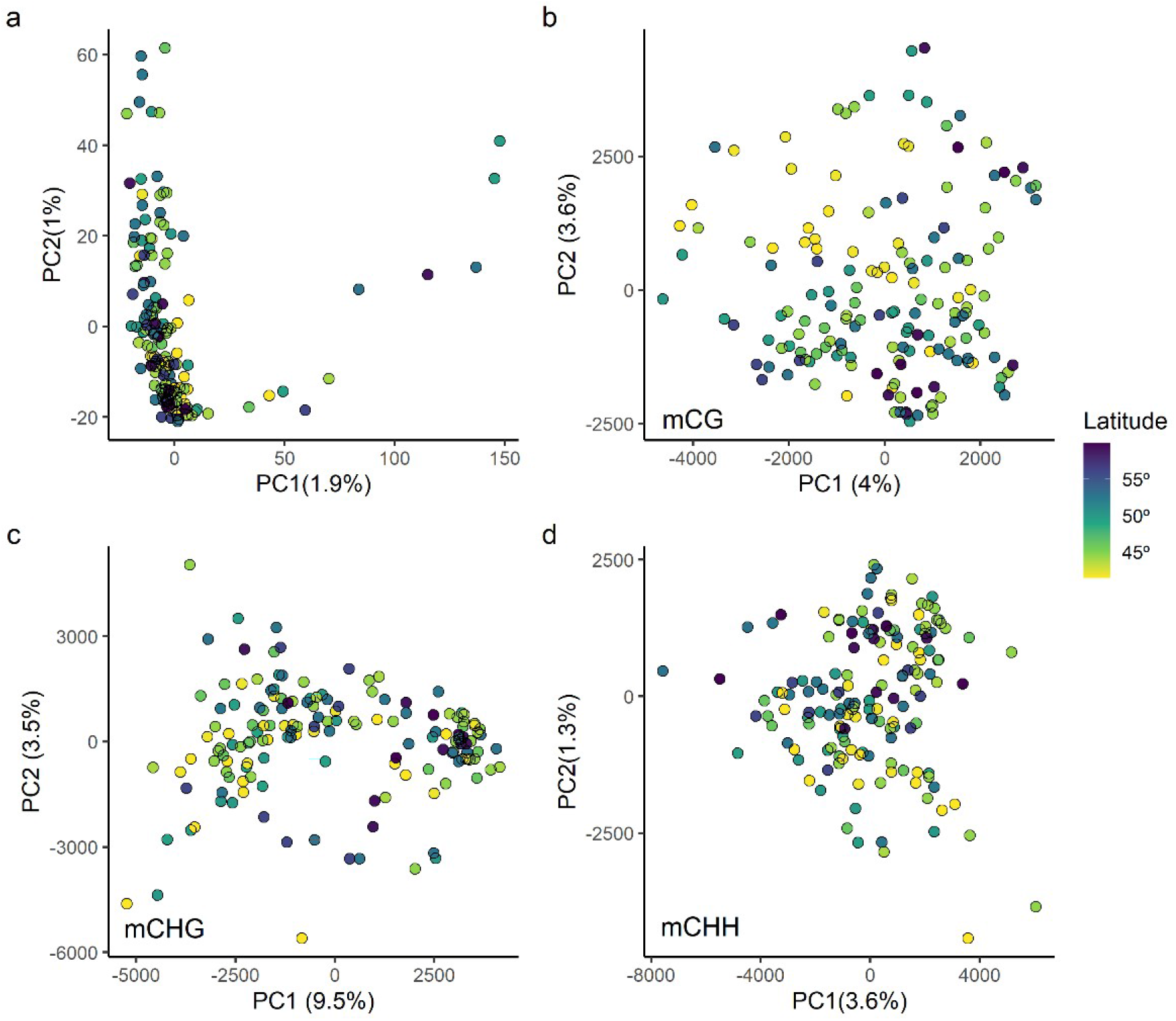
Genetic and epigenetic structure of the poplar ramets, colored according to latitude of origin. **a:** Genetic structure based on the SNPs called from WGBS data. **b-d**: Epigenetic structure for the CG (b), CHG (c) and CHH (d) sequence contexts.

### Genetic and epigenetic structure

To investigate a potential relationship between genetic and epigenetic structure in the Lombardy poplar, we conducted a Principal Component Analysis (PCA) based on methylated positions (MPs) and SNPs inferred from WGBS of the ramet samples. Among the sequenced ‘Italica’ clones, we did not find any clear genetic structure that could be associated with the geographic origin of the ramets (Figure 2a). As explained in Díez Rodríguez et al. (2022), the ramets that belonged to the ‘Italica’ cluster had a mean number of pairwise differences among individual ramets of around 96 SNPs out of the 4.906 investigated remaining positions. We targeted 4,906 loci equally distributed across the 19 *P. nigra* chromosomes selected from a larger set identified in Scaglione et. al (2019), which should allow for accurate and effective genotyping of population groups. To further assess if the loci targeted were actually sufficient for genotyping the populations analyzed, we called SNPs from the WGBS data. In this way, we increased the number of SNPs available for the study to 986,948 SNPs, mostly reflecting heterozygosity of the clonal genotype, not genetic differences between samples. After we removed SNPs heterozygous in > 95% of the samples and performed the pruning step, 343,977 SNPs remained for the analysis. Still, we did not find any genetic structure that could be associated with geographic patterns. On the other hand, despite the lack of genetic structure, some individuals with the same site of origin seemed to group together (Figures 2b and S5), indicating similar methylation profiles, specially in the CG context. Furthermore, when running PCA with MPs inside CDS (Figure S4), we observed some grouping, but this was not explained by any of the environmental variables that we tested (such as habitat type, elevation or habitat disturbance level).

### Relationship between methylation, geographic origin, and climate

To assess if there was any relationship between epigenetic variation, genetic variation, geographic origin, and climatic conditions, we analyzed the correlation between epigenetic distance and genetic, geographic, and climatic distance in ramets using mantel tests. We first correlated geographic with climatic distance, and genetic with both geographic and climatic distance. We found that climatic distance correlated with geographic distance (R = 0.7, p = 0.001), but genetic distance was not correlated with geographic distance or climatic distance (R = −0.03, p = ns, in both tests). We created epigenetic distance matrices based on MPs and DMRs. We did not find any correlation between epigenetic and genetic distance in any case, except for the MPs in the CHH context (Table 2). However, epigenetic distance significantly correlated with geographic and climatic distance in almost all cases. The highest correlation coefficients were found in the CG context between DMR-based epigenetic distance and both geographic and climatic distance (R=0.164 and p < 0.001, and R=0.141 and p < 0.001, respectively). Because geographic distance and climatic distance were strongly correlated, we ran partial mantel tests between epigenetic distance and climatic distance accounting for the geographic distance. In this case, most of the significant correlations disappeared, except for MPs in the CHH context.

**Table 2.**
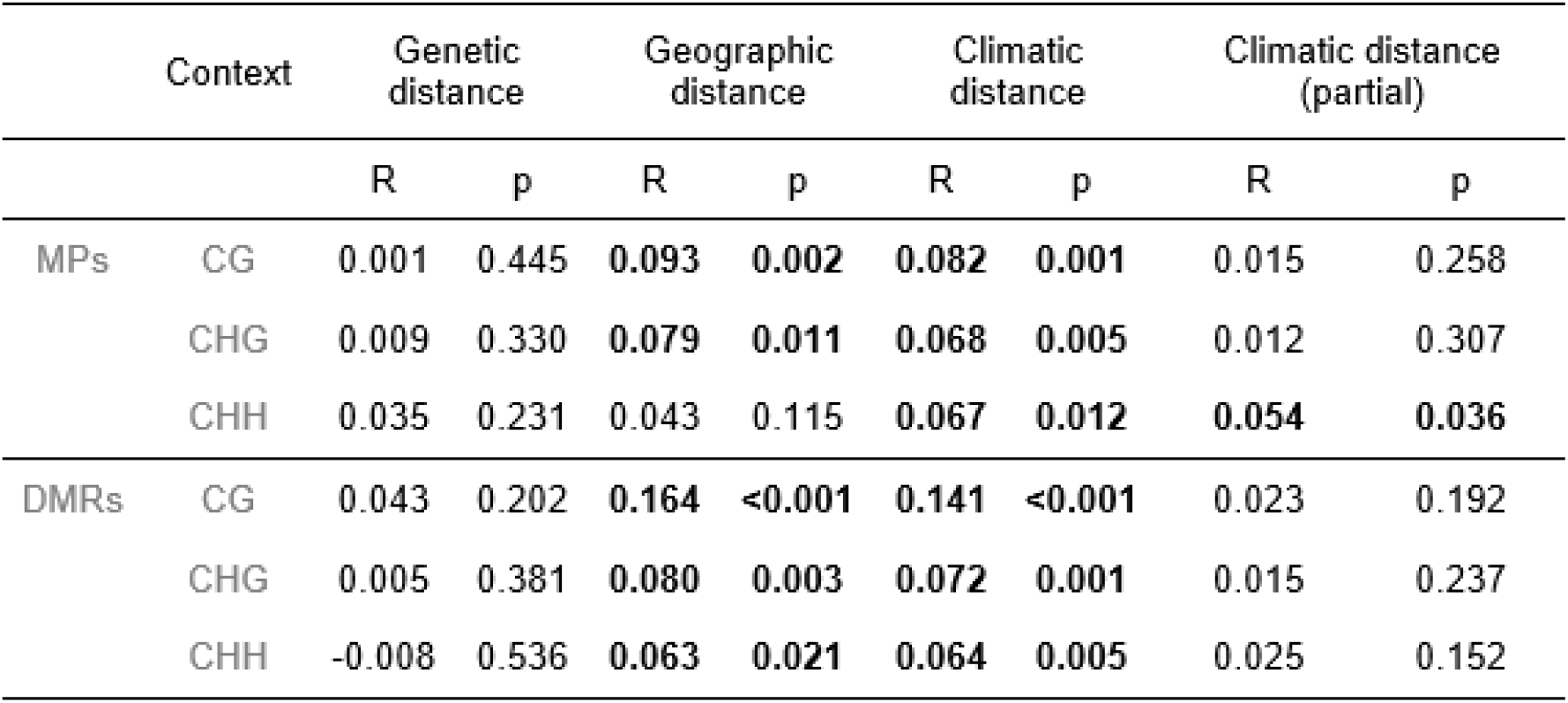
Mantel test coefficients for the correlation between epigenetic distance and genetic, geographic, and climatic distance in ramets. Epigenetic distance was tested both as individual methylated positions (MPs) and differentially methylated regions (DMRs). Significant correlations are highlighted in bold font.

To study the association between methylation patterns and climate of origin in more detail, we conducted a correlation analysis between global methylation levels in specific genomic features (i.e., promoters, coding sequences (CDS) and TEs) and bioclimatic variables (Figure 3). We found significant correlations in all sequence contexts, with the highest number of correlations observed in the CHH context. In fact, for the CHH context, we found correlations between all three genomic features and most temperature-related bioclimatic variables, such as maximum temperature and mean temperature related variables. Additionally, methylation levels in promoters and TEs in this context were negatively correlated with both latitude and longitude. On the other hand, methylation levels in the CG and CHG contexts showed no correlation with climatic variables, except for methylation in promoter and CDS regions and three precipitation variables (precipitation in the wettest month and wettest quarter, and precipitation in the warmest quarter). Furthermore, variables in CHH were grouped in a separate cluster while CG and CHG variables grouped mainly by genomic features.

**Figure 3.**
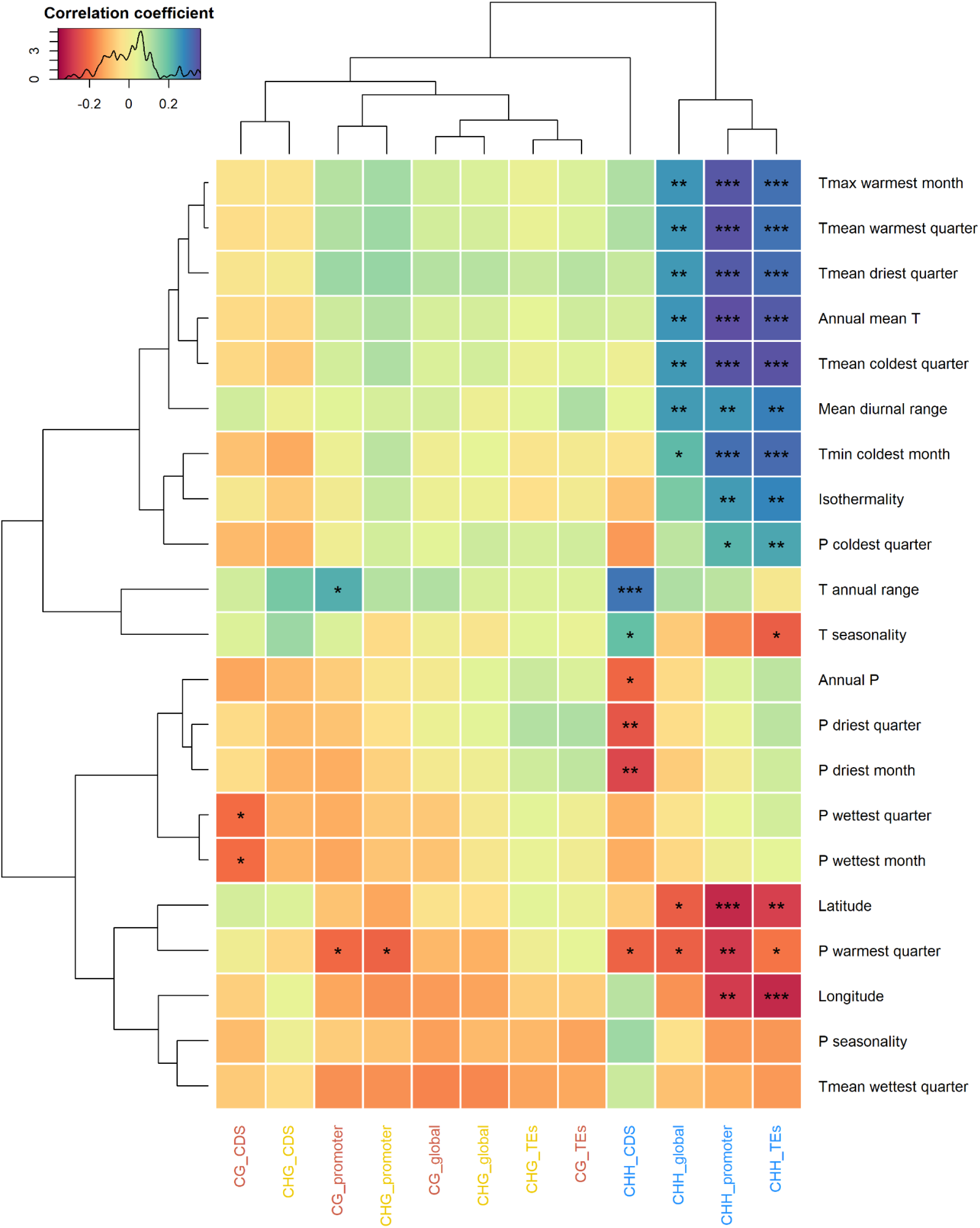
Spearman correlation analysis between global methylation levels in different genomic features and bioclimatic variables extracted from the CHELSA database. P = precipitation, T = temperature. P-values are adjusted for multiple pairwise comparisons using the “BH” method. Statistically significant correlations are labeled with the following code: p < 0.001 = ***; p < 0.01 = **; p < 0.05 = *. Variables are grouped by hierarchical clustering based on correlation coefficients.

### Persistence of DNA methylation patterns across clonal generations

To investigate if methylation patterns can be transmitted to the next clonal generation, we first compared average global methylation levels between ortets (parental individuals) and ramets (clonal offspring). In the ortets, methylation levels were consistently higher in all contexts (Figure 4A). The difference in global methylation levels between ortets and ramets was further evidenced by the number of hypermethylated ortet-vs-ramet DMRs (Figure 4B). When comparing ortets with their ramets, the number of DMRs in the CG context was considerably low for some groups (e.g. ES, IT2, FR1, CZ, NO), and the lowest of all contexts (10,180 total DMRs vs. 31,600 and 13,601 for CHG and CHH, respectively). On the other hand, the number of DMRs in the CHG and CHH contexts was more variable among different sites. Additionally, we conducted a PCA analysis using the paired clones (Figure S6) and found that pairs tended to group together, especially in the CG context.

**Figure 4.**
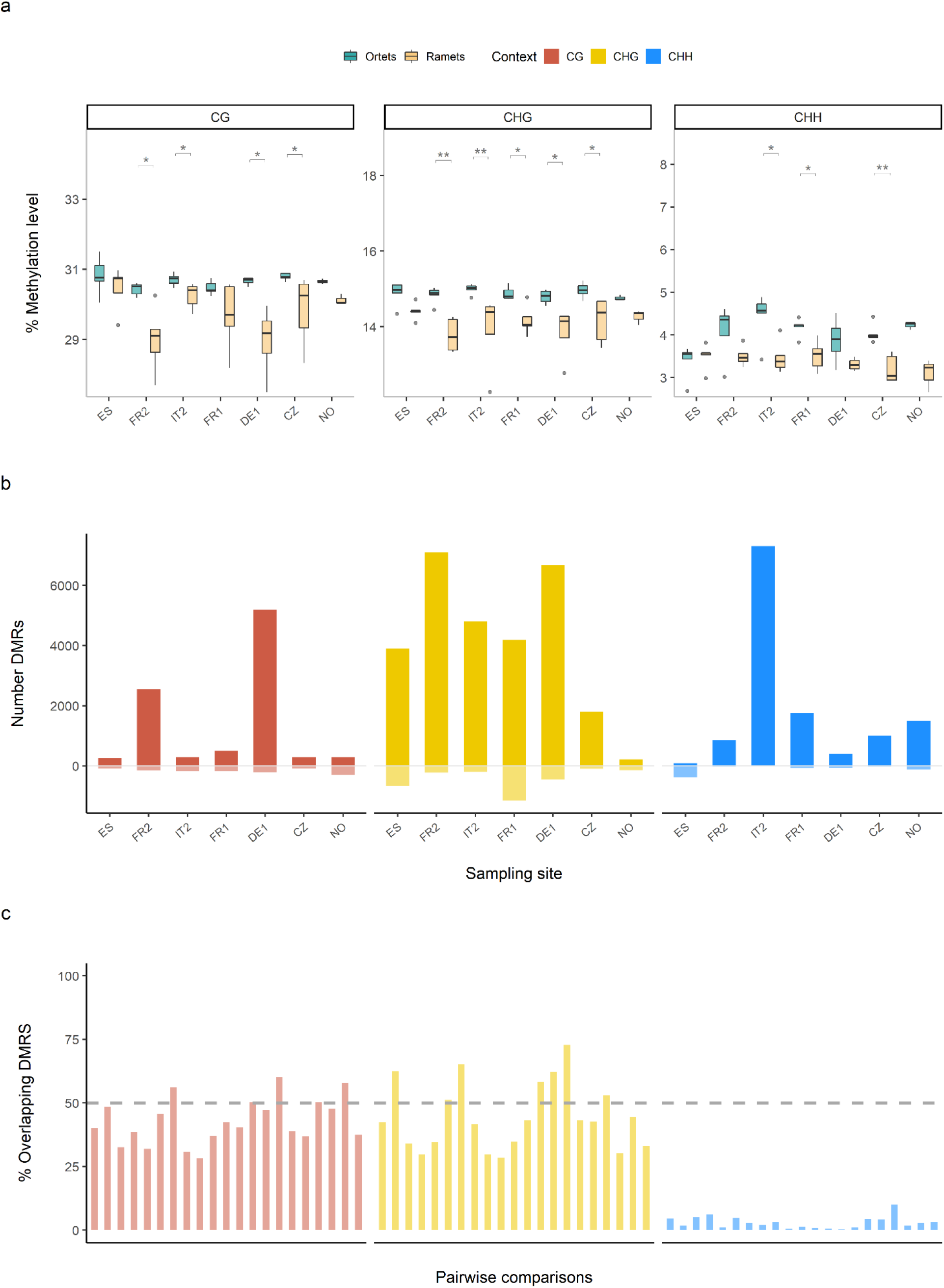
Differences in methylation profiles between ortets and ramets. a. Differences in global methylation levels between ortets (green) and their paired ramets (orange), for each sequence context. Statistically significant correlations are labeled with the following code: p < 0.001 = ***; p < 0.01 = **; p < 0.05 = *. P values were adjusted for multiple pairwise testing using the “BH” method. b. Total number of hypermethylated (above the 0 line) and hypomethylated (below the 0 line) DMRs between ortets and their paired ramets. c. Percentage of DMRs among ramet pairwise comparisons that overlap with DMRs among ortet pairwise comparisons. Each bar represents a pairwise comparison between ortets from each sampled site in Europe and the ramets of the same individuals. The dashed line indicates the threshold for 50% of ramet DMRs that overlap with ortet DMRs.

To further assess if methylation patterns persisted across clonal generations, we then intersected the DMRs found between pairwise comparisons in the ortets and the DMRs found between the ramets (Figure 4C). Between 25% and 50% of the ortet DMRs in CG and CHG overlapped with ramet DMRs. This percentage was considerably lower in the case of the CHH context, where less than 10% of the DMRs were also found in the ramets.

## Discussion

So far only few studies have used epigenomics to investigate the effects of environmentally induced epigenetic variation at a landscape level. Here, we present the first landscape-scale investigation of DNA methylation patterns in a system that has been almost exclusively clonally propagated. We found that average methylation levels were significantly correlated with climatic variables, specifically in TEs and gene promoter regions, and persisted across at least one clonal generation, despite the lack of evident genetic or epigenetic structure.

The lack of genetic structure can be explained by the very low genetic diversity found by genotyping the poplar clones (ramets) established in our common garden (Figure 2a). This was expected, given the clonal history of the ‘Italica’ cultivar. The ‘Italica’ cultivar likely originated from a single male clone in Central Asia, from where it spread to Europe. It is widely accepted that this clone was further artificially propagated from an individual or group of individuals found in Lombardy, Italy (Elwes and Henry, 1913). Our results suggest that a major fraction of the clones across Europe do indeed share a common line.

In a similar fashion, we did not find any clear epigenetic population structure but there appears to be some grouping in the CG context (Figure 2b) and epigenetic distance was positively correlated with geographic distance (Table 2). Furthermore, MPs inside CDS regions do show a pattern, but it was not explained by any of the environmental variables used in the analysis. This evidence points to the importance of other sources of epigenetic variation, such as genetic somatic mutations or stochastic epimutations. Several studies have reported age-related changes in the levels of cytosine methylation due to spontaneous methylation changes (Fraga *et al*., 2002; Dubrovina & Kiselev, 2016). Furthermore, Hofmeister *et al.* (2020) found evidence that spontaneous methylation changes are cumulative across somatic development in the close relative *Populus trichocarpa,* and that they have a higher rate than genetic mutations. Considering that the ‘Italica’ cultivar has been artificially propagated for the last two centuries, stochastic epimutations have likely accumulated across several clonal generations, confounding any environmentally induced epigenetic population structure.

Previous studies on population epigenomics have found that epigenetic variation is associated with genetic variation in Brassicaceae (Dubin *et al,* 2015, Kawakatsu *et al*., 2016; Galanti *et al*., 2022), thus hindering the study of the relationship between environmental epigenetic variation and climatic conditions. The use of a clonal cultivar circumvents this problem. We used mantel tests to investigate if epigenetic distance, measured as the distance between both single methylated variants (MPs) and differentially methylated regions (DMRs), was correlated with genetic, geographic and/or climatic distance (Table 2). We found that epigenetic distance did not correlate with genetic distance in all cases except one (MPs in the CHH context) but correlated with both geographic distance and climatic distance in almost all cases (see also Figure S2). However, when accounting for geographic distance, the correlations with climatic distance disappeared, except for MPs in the CHH context. As suggested above, if stochastic epimutations are contributing to a major fraction of the epigenetic variation, the correlation between epigenetic distance and geographic distance could be explained by isolation-by-distance processes, since this cultivar was gradually propagated across Europe (Slatkin, 1993). This evidence thus suggests that epigenetic variation of the individuals analyzed might be both under environmental and stochastic control.

To assess whether the methylation profiles under climatic control could potentially have a functional role, we extracted the methylation levels of specific genomic features (gene promoters, gene body and transposable elements, specifically). We then correlated methylation levels with individual bioclimatic variables (Figure 3). Methylation levels were strongly correlated with most temperature variables, particularly in the case of gene promoters and TEs in the CHH context, which would also explain the correlation with latitude and longitude. Our results are in line with previous studies that have reported the potential effects of temperature on DNA methylation in several plant organisms (Dubin et al., 2015; Conde et al., 2017; Zhang et al., 2018; Galanti et al., 2022; Sammarco et al., 2022). On the other hand, methylation levels correlated with very few precipitation variables but, as opposed to temperature variables, we observed more significant correlations in the CG and CHG context. It is conceivable that a certain degree of environmental information regarding water availability might be encoded in more stable methylation contexts and transmitted to the clonal offspring, since *Populus nigra* is a riparian species that depends on river flooding regimes for successful seed and cutting dispersal (Smulders *et al*., 2008). Nevertheless, our results indicate that methylation patterns in CHH might be highly dynamic and rapidly respond to new environmental cues. This assumption is further supported by the changes in global methylation levels observed between ortet-ramet pairs (Figure 4A). Although there were almost no differences in methylation levels between individuals from different geographic origins in any of the contexts, methylation levels were significantly higher in the ortets than in the corresponding ramets for many locations. In poplar, methylation levels have been shown to increase under drought conditions (Raj *et al*., 2011; Peña Pontón *et al,* 2022). Given that 2018 was a year characterized by particularly extreme drought events in Europe, and the ramets were well watered during the whole summer, it is possible that the differences in methylation levels between ortets and ramets are the result of differences in water availability. Furthermore, we observed a considerable decrease in the number of DMRs found among ramets (Supplementary Figure 2), suggesting that methylation profiles in leaves in the CHH context might have already adjusted to the new conditions of the common garden.

Despite these dynamic changes in CHH methylation, a considerable fraction of the methylation patterns appeared to be transmitted to the clonal offspring, particularly in the CG and CHG contexts. We found that approximately 25% of the DMRs in CG and CHG called from pairwise comparisons among the ramets of different sampled sites overlapped with the DMRs found among the ramets of the same pairwise comparison (Figure 4C). The fact that we could find these specific regions both in the ortets and the ramets provides further evidence that methylation patterns in the CG and CHG contexts can potentially be transmitted to the clonal offspring. Conversely, less than 10% of the DMRs found in the CHH context were transmitted to the next clonal generation. This further supports our conclusion that methylation in the CHH context is highly dynamic. It is, however, challenging to determine if there was an active change in the methylome as a result of new environmental cues, or if these patterns are established *de novo* every year in leaf tissue. If in fact leaf CHH methylation patterns are determined in every new season, this could possibly explain the low number of DMRs observed in the CHH context, both among the ortets and the ramets (Supplementary Figure S2). If the environmental conditions in the common garden resemble those of the original sites, then the methylome in CHH in the ramets would also resemble the methylome of the ortets. If the conditions are nothing alike, then a higher number of DMRs would be expected. Based on the total number of DMRs, the latter might be true. The number of DMRs was considerably higher when comparing ortets sampled in Spain with ortets sampled in Northern European sites (Figure S2), while only a few DMRs were found between sites that belong to similar Köppen climatic areas (e.g., FR1 vs FR2). In the common garden, however, where the environmental conditions were the same for all the individuals, the number of total DMRs between ramets from different sites was very low, suggesting that the ramets might have rapidly adjusted to common garden conditions. As proposed by Ito and colleagues (2019), DNA methylation in natural environments might have two components, genomic regions that might change dynamically and epigenetic marks for stable gene expression that are rather fixed. If this is the case, it opens interesting new research possibilities, if a certain fraction of epigenetic information is stored in symmetrical stable contexts, but some of it can rapidly shift to reflect new environments. In practical terms, this would imply that methylation variation is partitioned in distinct “modules”, and further experiments should target individual sources of environmentally induced epigenetic variation.

In summary, our study is the first landscape-scale investigation of DNA methylation patterns in a system that has been almost exclusively clonally propagated. We found that methylation patterns in the Lombardy poplar are independent of genetic structure, but that methylation profiles are associated with climatic conditions. Furthermore, we have shown that a fraction of DMRs is transmitted to the next clonal generation, and that methylation in the CHH levels is highly dynamic and might rapidly adjust to new environmental conditions. Our results suggest that the CHH context is the most responsive to changing environments and that the stability of induced changes across clonal generations is stronger in CG and CHG. We have shown that the Lombardy poplar is a valuable system to study environmentally induced epigenetic variation in a naturally occurring near-isogenic population, with limited confounding genetic variation. Our study provides further insight into how methylation patterns in natural populations might vary along geographic and climatic gradients. However, further research is necessary to assess whether DNA methylation can have an effect on phenotypic plasticity. The high resolution methylome data generated in our experiment is a significant resource for Epigenome Wide Association Studies (EWAS), and can considerably contribute to our understanding of how methylation variation affects plant acclimation and adaptation.

## Acknowledgements

We thank members of the EpiDiverse consortium (www.epidiverse.eu) for valuable inputs during preparation and execution of the study, and reviewers for useful comments. For computing, we acknowledge Prof. Peter Stadler at the University of Leipzig and David Langenberger from ecSeq, for hosting the EpiDiverse servers.

## Author contributions

**Bárbara Díez Rodríguez:** Methodology, formal analysis, Investigation, Software - implementation of the computer code and supporting algorithms; testing of existing code components, writing - original draft, visualization

**Dario Galanti**: Formal analysis (WGBS SNPs), Software - implementation of the computer code and supporting algorithms; testing of existing code components; Writing – Review & Editing

**Adam Nunn:** Software – Programming, software development, implementation of the computer code and supporting algorithms; testing of existing code components; Data curation, Writing – Review & Editing

**Cristian Peña:** Software - implementation of the computer code and supporting algorithms; testing of existing code components, Writing – Review & Editing

**Paloma Pérez-Bello:** Software - implementation of the computer code and supporting algorithms; testing of existing code components, Writing – Review & Editing

**Iris Sammarco**: Software - implementation of the computer code and supporting algorithms; testing of existing code components; Writing – Review & Editing

**Claude Becker:** Writing – Review & Editing, Supervision

**Katharina Jandrasits:** Investigation – Library preparation

**Emanuele de Paoli:** Writing – Review & Editing

**Koen J.F. Verhoeven**: Writing – Review & Editing, Supervision, Project administration, Funding acquisition

**Lars Opgenoorth:** Conceptualization, Writing – Review & Editing, Supervision, Project administration, Funding acquisition

**Katrin Heer:** Conceptualization, Writing – Review & Editing, Supervision, Project administration, Funding acquisition

## Data availability

The *Populus nigra* cv ‘Italica’ reference genome and the genome annotations are freely available at the European Nucleotide Archive (ENA) under project number PRJEB44889. All sequenced raw fastq files are available under project number PRJEB44879. Bioclimatic data for all sequenced individuals is available in Zenodo at https://doi.org/10.5281/zenodo.5995424.

