## SupplementaryMaterial for "Epigenetic variation in the Lombardy poplar along climatic gradients is independent of genetic structure and persists across clonal reproduction"

### Supplementary information

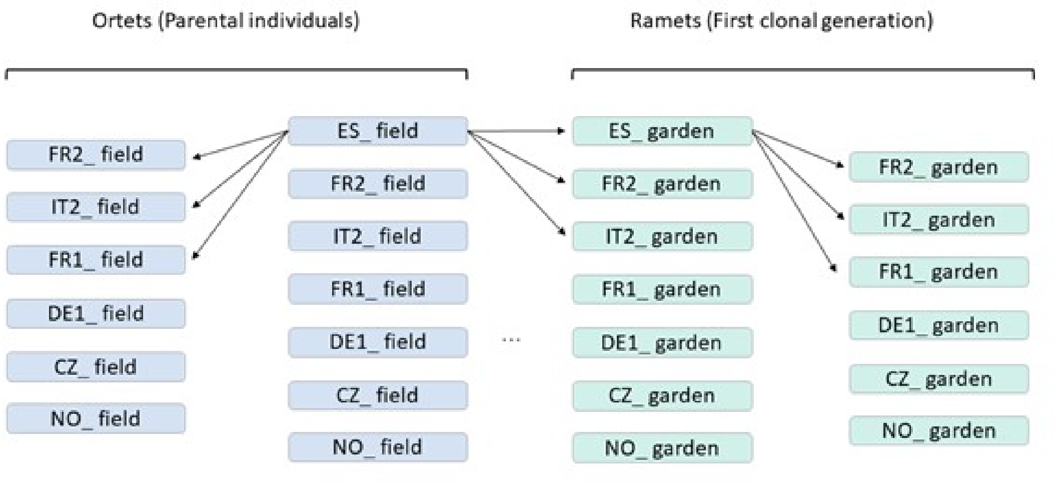


Supplementary Figure 1. Multiple pairwise comparison design for DMR calling groups. DMRs were called in three different instances. First, we run the pipeline using only groups containing ortet (field) samples in each pairwise comparison; second, we compared groups containing only ramet (garden) samples, and third, we compared ortet samples with their paired ramet samples (field vs garden).

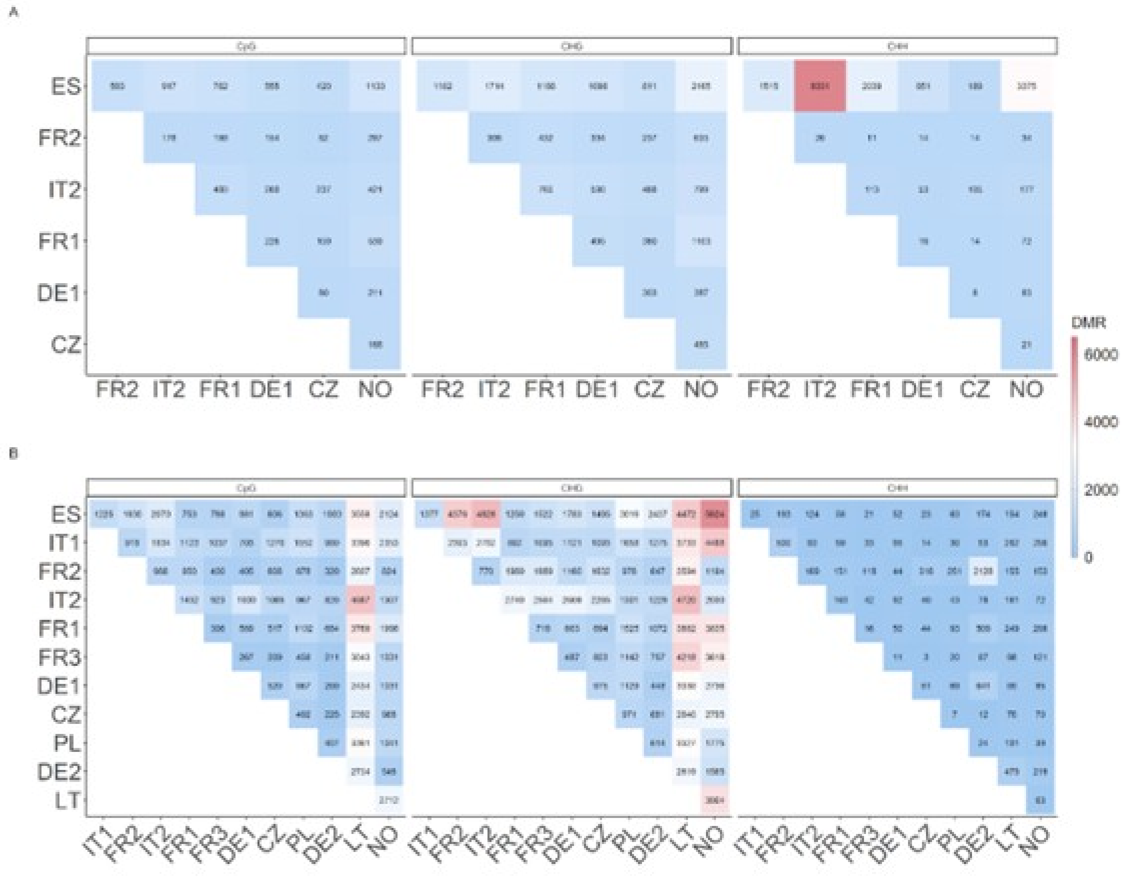


Supplementary Figure 2. Total number of DMRs between pairwise comparisons for A) ortets and B) ramets. Sampling sites are ordered from South to North, according to latitude.

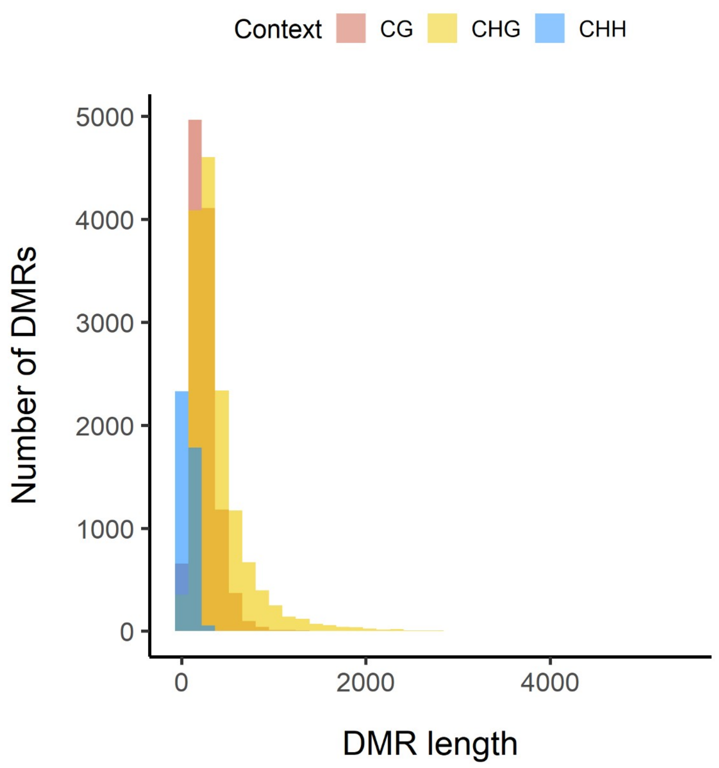


Supplementary figure 3. Histogram of the length of merged DMRs for each context.

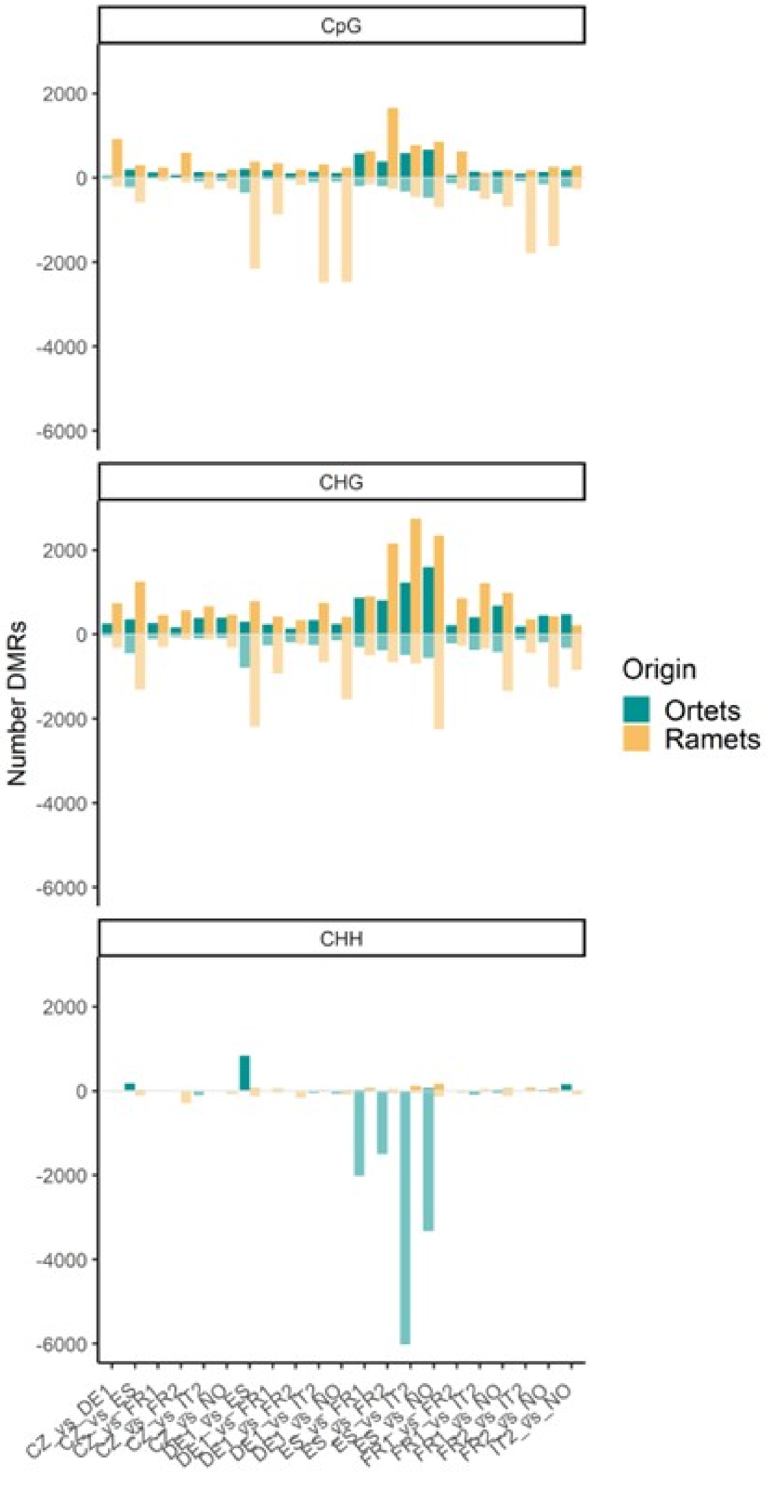


Supplementary Figure 4. Total number of hyper (above the 0 line) and hypomethylated (below the 0 line) DMRs in all pairwise comparisons among paired ortets (dark green) and ramets (orange).

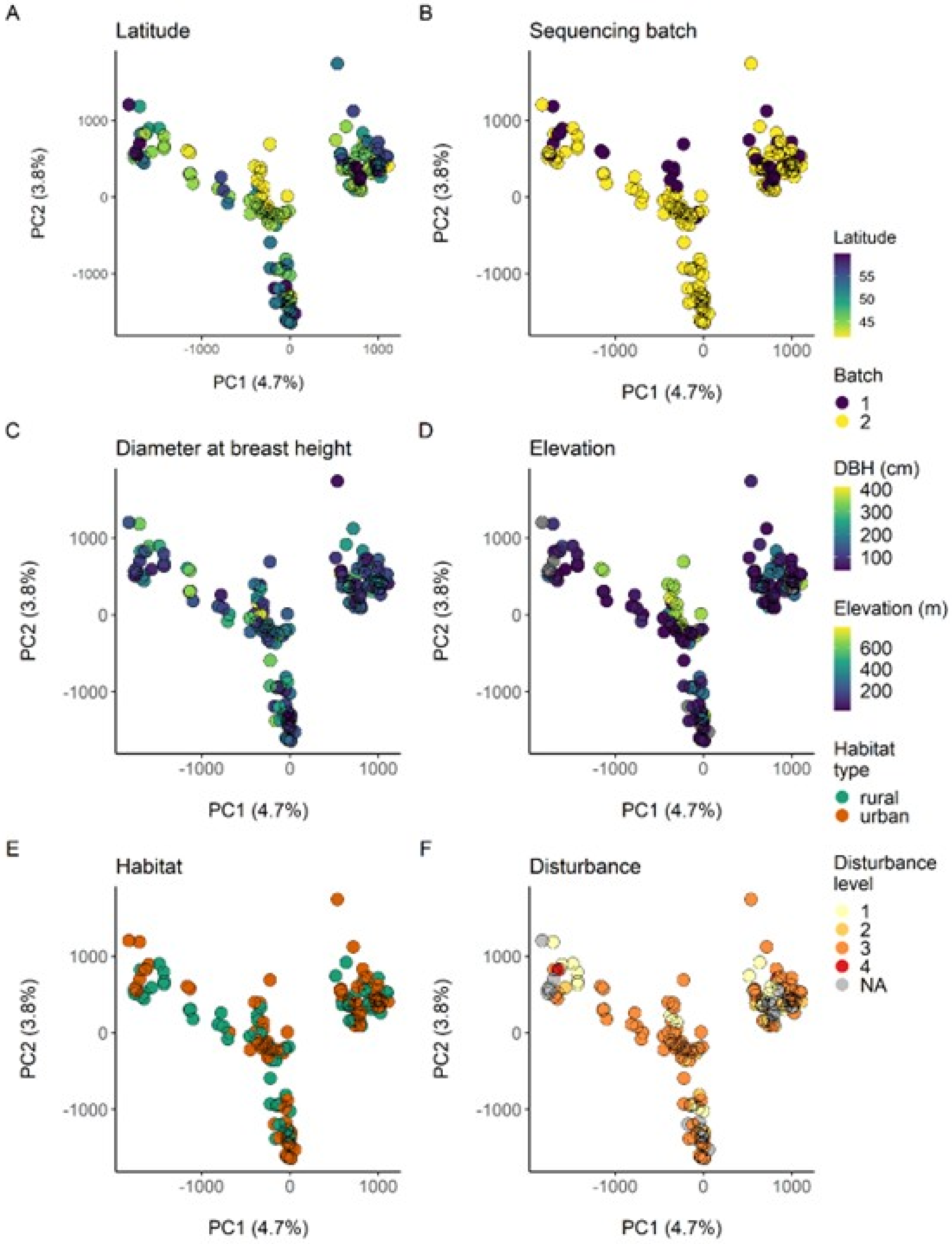


Supplementary Figure 5. Epigenetic structure using only methylated positions found inside coding sequences in the CG context, colored according to different environmental variables.

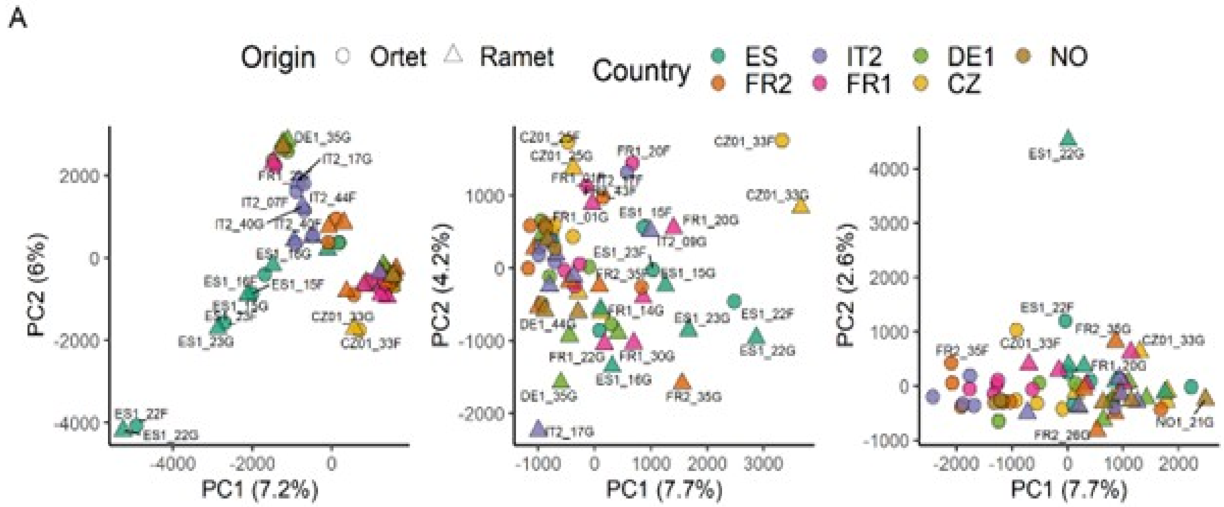


Supplementary Figure 6. PCAs of epigenetic structure using methylated positions inside coding regions (CDS), comparing ortets and paired ramets, for CG, CHG and CHH contexts.

Supplementary table 1. Mapping stats and bisulfite non-conversion rate for each sequenced individual sample.

| SampleID | Site | Origin | Coverage | Mapping Rate (%) | Bisulfite NonConv Rate (%) |
|---|---|---|---|---|---|
| PN_CZ_01_01_R1_GG0 | C.Republic | Ramets | 52,88 | 80,22 | 0,005 |
| PN_CZ_01_03_R1_GG0 | C.Republic | Ramets | 42,13 | 80,99 | 0,005 |
| PN_CZ_01_13_R1_GG0 | C.Republic | Ramets | 38,88 | 80,00 | 0,005 |
| PN_CZ_01_16_R1_GG0 | C.Republic | Ramets | 53,44 | 79,62 | 0,005 |
| PN_CZ_01_20_R1_GG0 | C.Republic | Ramets | 44,50 | 80,26 | 0,005 |
| PN_CZ_01_22_R1_GG0 | C.Republic | Ramets | 72,50 | 78,60 | 0,004 |
| PN_CZ_01_25_R1_GG0 | C.Republic | Ramets | 91,00 | 80,47 | 0,005 |
| PN_CZ_01_29_R1_GG0 | C.Republic | Ramets | 38,50 | 79,51 | 0,005 |
| PN_CZ_01_33_R1_GG0 | C.Republic | Ramets | 81,00 | 79,67 | 0,004 |
| PN_CZ_01_37_R1_GG0 | C.Republic | Ramets | 39,94 | 79,65 | 0,005 |
| PN_CZ_01_39_R1_GG0 | C.Republic | Ramets | 50,81 | 78,83 | 0,005 |
| PN_CZ_01_40_R1_GG0 | C.Republic | Ramets | 64,13 | 79,79 | 0,006 |
| PN_CZ_01_44_R1_GG0 | C.Republic | Ramets | 43,38 | 80,36 | 0,005 |
| PN_CZ_01_52_R1_GG0 | C.Republic | Ramets | 61,19 | 80,17 | 0,005 |
| PN_DE_01_06_R1_GG0 | Germany1 | Ramets | 47,63 | 80,10 | 0,004 |
| PN_DE_01_08_R1_GG0 | Germany1 | Ramets | 46,00 | 78,70 | 0,004 |
| PN_DE_01_10_R1_GG0 | Germany1 | Ramets | 76,63 | 37,41 | 0,005 |
| PN_DE_01_14_R1_GG0 | Germany1 | Ramets | 57,94 | 78,89 | 0,004 |
| PN_DE_01_15_R1_GG0 | Germany1 | Ramets | 47,75 | 79,16 | 0,004 |
| PN_DE_01_22_R1_GG0 | Germany1 | Ramets | 63,25 | 80,14 | 0,005 |
| PN_DE_01_29_R1_GG0 | Germany1 | Ramets | 57,25 | 79,53 | 0,004 |
| PN_DE_01_31_R1_GG0 | Germany1 | Ramets | 53,19 | 79,60 | 0,006 |
| PN_DE_01_35_R1_GG0 | Germany1 | Ramets | 62,31 | 79,17 | 0,005 |

| | | | | | |
|---|---|---|---|---|---|
| PN_DE_01_38_R1_GG0 | Germany1 | Ramets | 53,56 | 78,74 | 0,008 |
| PN_DE_01_41_R1_GG0 | Germany1 | Ramets | 57,31 | 79,83 | 0,005 |
| PN_DE_01_44_R1_GG0 | Germany1 | Ramets | 52,81 | 78,38 | 0,006 |
| PN_DE_01_53_R1_GG0 | Germany1 | Ramets | 50,31 | 80,01 | 0,005 |
| PN_DE_02_02_R1_GG0 | Germany2 | Ramets | 40,75 | 79,45 | 0,005 |
| PN_DE_02_08_R1_GG0 | Germany2 | Ramets | 44,56 | 79,58 | 0,006 |
| PN_DE_02_10_R1_GG0 | Germany2 | Ramets | 51,38 | 80,64 | 0,005 |
| PN_DE_02_13_R1_GG0 | Germany2 | Ramets | 75,56 | 79,56 | 0,005 |
| PN_DE_02_15_R1_GG0 | Germany2 | Ramets | 45,13 | 80,63 | 0,006 |
| PN_DE_02_19_R1_GG0 | Germany2 | Ramets | 51,25 | 79,58 | 0,005 |
| PN_DE_02_20_R1_GG0 | Germany2 | Ramets | 58,19 | 79,02 | 0,005 |
| PN_DE_02_24_R1_GG0 | Germany2 | Ramets | 41,81 | 78,63 | 0,005 |
| PN_DE_02_33_R1_GG0 | Germany2 | Ramets | 50,94 | 79,64 | 0,005 |
| PN_DE_02_38_R1_GG0 | Germany2 | Ramets | 46,44 | 78,39 | 0,005 |
| PN_DE_02_40_R1_GG0 | Germany2 | Ramets | 64,81 | 79,60 | 0,004 |
| PN_DE_02_41_R1_GG0 | Germany2 | Ramets | 53,81 | 80,39 | 0,005 |
| PN_DE_02_50_R1_GG0 | Germany2 | Ramets | 46,31 | 79,50 | 0,005 |
| PN_DE_02_53_R1_GG0 | Germany2 | Ramets | 68,94 | 79,46 | 0,005 |
| PN_ES_01_01_R1_GG0 | Spain | Ramets | 48,69 | 78,75 | 0,004 |
| PN_ES_01_04_R1_GG0 | Spain | Ramets | 42,06 | 79,40 | 0,005 |
| PN_ES_01_05_R1_GG0 | Spain | Ramets | 41,56 | 78,88 | 0,005 |
| PN_ES_01_06_R1_GG0 | Spain | Ramets | 33,19 | 78,69 | 0,005 |
| PN_ES_01_07_R1_GG0 | Spain | Ramets | 53,19 | 78,03 | 0,005 |
| PN_ES_01_08_R1_GG0 | Spain | Ramets | 44,13 | 78,12 | 0,005 |
| PN_ES_01_09_R1_GG0 | Spain | Ramets | 49,50 | 77,55 | 0,005 |
| PN_ES_01_10_R1_GG0 | Spain | Ramets | 46,06 | 77,78 | 0,006 |
| PN_ES_01_11_R1_GG0 | Spain | Ramets | 50,38 | 78,02 | 0,005 |
| PN_ES_01_15_R1_GG0 | Spain | Ramets | 53,06 | 78,21 | 0,005 |

| | | | | | |
|---|---|---|---|---|---|
| PN_ES_01_16_R1_GG0 | Spain | Ramets | 55,00 | 78,71 | 0,005 |
| PN_ES_01_17_R1_GG0 | Spain | Ramets | 56,06 | 77,21 | 0,005 |
| PN_ES_01_22_R1_GG0 | Spain | Ramets | 45,19 | 77,17 | 0,006 |
| PN_ES_01_23_R1_GG0 | Spain | Ramets | 67,38 | 80,00 | 0,004 |
| PN_FR_01_01_R1_GG0 | France1 | Ramets | 42,06 | 80,10 | 0,005 |
| PN_FR_01_08_R1_GG0 | France1 | Ramets | 37,63 | 80,22 | 0,004 |
| PN_FR_01_09_R1_GG0 | France1 | Ramets | 35,94 | 79,73 | 0,004 |
| PN_FR_01_14_R1_GG0 | France1 | Ramets | 43,69 | 78,97 | 0,004 |
| PN_FR_01_20_R1_GG0 | France1 | Ramets | 42,19 | 80,86 | 0,004 |
| PN_FR_01_21_R1_GG0 | France1 | Ramets | 37,44 | 78,86 | 0,005 |
| PN_FR_01_22_R1_GG0 | France1 | Ramets | 148,69 | 78,82 | 0,004 |
| PN_FR_01_24_R1_GG0 | France1 | Ramets | 59,69 | 78,47 | 0,004 |
| PN_FR_01_30_R1_GG0 | France1 | Ramets | 43,38 | 77,88 | 0,004 |
| PN_FR_01_34_R1_GG0 | France1 | Ramets | 60,75 | 78,71 | 0,005 |
| PN_FR_01_44_R1_GG0 | France1 | Ramets | 46,38 | 79,58 | 0,005 |
| PN_FR_01_50_R1_GG0 | France1 | Ramets | 46,75 | 79,98 | 0,005 |
| PN_FR_01_51_R1_GG0 | France1 | Ramets | 54,00 | 79,04 | 0,004 |
| PN_FR_01_57_R1_GG0 | France1 | Ramets | 62,63 | 79,27 | 0,004 |
| PN_FR_02_07_R1_GG0 | France2 | Ramets | 61,00 | 79,85 | 0,005 |
| PN_FR_02_09_R1_GG0 | France2 | Ramets | 70,19 | 76,80 | 0,01 |
| PN_FR_02_12_R1_GG0 | France2 | Ramets | 59,94 | 78,61 | 0,005 |
| PN_FR_02_14_R1_GG0 | France2 | Ramets | 47,38 | 77,60 | 0,005 |
| PN_FR_02_15_R1_GG0 | France2 | Ramets | 58,19 | 78,09 | 0,005 |
| PN_FR_02_20_R1_GG0 | France2 | Ramets | 53,88 | 77,10 | 0,005 |
| PN_FR_02_26_R1_GG0 | France2 | Ramets | 40,31 | 78,73 | 0,005 |
| PN_FR_02_35_R1_GG0 | France2 | Ramets | 41,88 | 78,38 | 0,005 |
| PN_FR_02_37_R1_GG0 | France2 | Ramets | 43,38 | 78,13 | 0,005 |
| PN_FR_02_39_R1_GG0 | France2 | Ramets | 68,31 | 79,37 | 0,004 |

| | | | | | |
|---|---|---|---|---|---|
| PN_FR_02_41_R1_GG0 | France2 | Ramets | 61,13 | 80,25 | 0,005 |
| PN_FR_02_43_R1_GG0 | France2 | Ramets | 60,75 | 80,06 | 0,006 |
| PN_FR_02_49_R1_GG0 | France2 | Ramets | 46,19 | 78,94 | 0,004 |
| PN_FR_03_03_R1_GG0 | France3 | Ramets | 48,56 | 79,82 | 0,004 |
| PN_FR_03_06_R1_GG0 | France3 | Ramets | 43,25 | 77,59 | 0,004 |
| PN_FR_03_08_R1_GG0 | France3 | Ramets | 73,63 | 79,34 | 0,005 |
| PN_FR_03_10_R1_GG0 | France3 | Ramets | 69,13 | 79,97 | 0,004 |
| PN_FR_03_13_R1_GG0 | France3 | Ramets | 66,69 | 79,54 | 0,004 |
| PN_FR_03_15_R1_GG0 | France3 | Ramets | 54,56 | 80,12 | 0,004 |
| PN_FR_03_18_R1_GG0 | France3 | Ramets | 52,50 | 80,39 | 0,005 |
| PN_FR_03_20_R1_GG0 | France3 | Ramets | 49,81 | 80,39 | 0,004 |
| PN_FR_03_26_R1_GG0 | France3 | Ramets | 52,50 | 80,94 | 0,005 |
| PN_FR_03_31_R1_GG0 | France3 | Ramets | 33,63 | 78,94 | 0,004 |
| PN_FR_03_35_R1_GG0 | France3 | Ramets | 38,13 | 79,73 | 0,004 |
| PN_FR_03_41_R1_GG0 | France3 | Ramets | 48,31 | 79,74 | 0,005 |
| PN_FR_03_53_R1_GG0 | France3 | Ramets | 91,13 | 79,71 | 0,004 |
| PN_FR_03_56_R1_GG0 | France3 | Ramets | 58,81 | 80,07 | 0,004 |
| PN_IT_01_06_R1_GG0 | Italy1 | Ramets | 60,69 | 79,54 | 0,005 |
| PN_IT_01_07_R1_GG0 | Italy1 | Ramets | 49,69 | 79,93 | 0,005 |
| PN_IT_01_08_R1_GG0 | Italy1 | Ramets | 46,81 | 80,25 | 0,005 |
| PN_IT_01_21_R1_GG0 | Italy1 | Ramets | 41,69 | 78,15 | 0,005 |
| PN_IT_01_25_R1_GG0 | Italy1 | Ramets | 57,50 | 77,76 | 0,005 |
| PN_IT_01_32_R1_GG0 | Italy1 | Ramets | 56,69 | 78,36 | 0,005 |
| PN_IT_01_33_R1_GG0 | Italy1 | Ramets | 63,50 | 78,09 | 0,005 |
| PN_IT_01_35_R1_GG0 | Italy1 | Ramets | 57,31 | 77,52 | 0,005 |
| PN_IT_01_38_R1_GG0 | Italy1 | Ramets | 54,75 | 77,75 | 0,005 |
| PN_IT_01_41_R1_GG0 | Italy1 | Ramets | 64,69 | 77,67 | 0,005 |
| PN_IT_01_43_R1_GG0 | Italy1 | Ramets | 47,00 | 79,55 | 0,005 |

| | | | | | |
|---|---|---|---|---|---|
| PN_IT_01_47_R1_GG0 | Italy1 | Ramets | 53,13 | 79,68 | 0,005 |
| PN_IT_01_50_R1_GG0 | Italy1 | Ramets | 60,69 | 78,86 | 0,004 |
| PN_IT_02_02_R1_GG0 | Italy2 | Ramets | 54,13 | 80,25 | 0,004 |
| PN_IT_02_05_R1_GG0 | Italy2 | Ramets | 48,19 | 80,46 | 0,005 |
| PN_IT_02_07_R1_GG0 | Italy2 | Ramets | 42,00 | 79,74 | 0,005 |
| PN_IT_02_09_R1_GG0 | Italy2 | Ramets | 61,63 | 79,84 | 0,005 |
| PN_IT_02_17_R1_GG0 | Italy2 | Ramets | 56,25 | 79,08 | 0,005 |
| PN_IT_02_19_R1_GG0 | Italy2 | Ramets | 46,56 | 79,10 | 0,005 |
| PN_IT_02_28_R1_GG0 | Italy2 | Ramets | 62,19 | 79,09 | 0,006 |
| PN_IT_02_31_R1_GG0 | Italy2 | Ramets | 59,19 | 79,45 | 0,006 |
| PN_IT_02_34_R1_GG0 | Italy2 | Ramets | 48,19 | 79,55 | 0,006 |
| PN_IT_02_35_R1_GG0 | Italy2 | Ramets | 48,56 | 79,64 | 0,006 |
| PN_IT_02_38_R1_GG0 | Italy2 | Ramets | 44,31 | 79,17 | 0,006 |
| PN_IT_02_40_R1_GG0 | Italy2 | Ramets | 44,56 | 79,78 | 0,006 |
| PN_IT_02_44_R1_GG0 | Italy2 | Ramets | 72,13 | 79,34 | 0,005 |
| PN_IT_02_48_R1_GG0 | Italy2 | Ramets | 36,13 | 80,08 | 0,004 |
| PN_LT_01_01_R1_GG0 | Lithuania | Ramets | 50,69 | 80,52 | 0,006 |
| PN_LT_01_03_R1_GG0 | Lithuania | Ramets | 68,25 | 79,98 | 0,005 |
| PN_LT_01_04_R1_GG0 | Lithuania | Ramets | 77,06 | 78,83 | 0,005 |
| PN_LT_01_06_R1_GG0 | Lithuania | Ramets | 43,44 | 80,24 | 0,005 |
| PN_LT_01_08_R1_GG0 | Lithuania | Ramets | 64,31 | 79,03 | 0,005 |
| PN_LT_01_09_R1_GG0 | Lithuania | Ramets | 44,38 | 79,66 | 0,005 |
| PN_LT_01_10_R1_GG0 | Lithuania | Ramets | 53,69 | 79,99 | 0,005 |
| PN_LT_01_11_R1_GG0 | Lithuania | Ramets | 56,00 | 79,85 | 0,005 |
| PN_LT_01_12_R1_GG0 | Lithuania | Ramets | 46,81 | 80,40 | 0,006 |
| PN_NO_01_11_R1_GG0 | Norway | Ramets | 53,69 | 79,42 | 0,004 |
| PN_NO_01_18_R1_GG0 | Norway | Ramets | 67,94 | 78,13 | 0,005 |
| PN_NO_01_19_R1_GG0 | Norway | Ramets | 57,31 | 78,44 | 0,005 |

| | | | | | |
|---|---|---|---|---|---|
| PN_NO_01_21_R1_GG0 | Norway | Ramets | 52,25 | 77,84 | 0,005 |
| PN_NO_01_22_R1_GG0 | Norway | Ramets | 52,00 | 78,35 | 0,005 |
| PN_NO_01_26_R1_GG0 | Norway | Ramets | 57,75 | 78,36 | 0,005 |
| PN_NO_01_33_R1_GG0 | Norway | Ramets | 51,56 | 76,95 | 0,006 |
| PN_NO_01_36_R1_GG0 | Norway | Ramets | 43,75 | 79,59 | 0,005 |
| PN_NO_01_38_R1_GG0 | Norway | Ramets | 44,88 | 79,95 | 0,005 |
| PN_NO_01_41_R1_GG0 | Norway | Ramets | 74,06 | 80,60 | 0,005 |
| PN_NO_01_49_R1_GG0 | Norway | Ramets | 50,31 | 80,40 | 0,005 |
| PN_NO_01_50_R1_GG0 | Norway | Ramets | 58,88 | 79,98 | 0,005 |
| PN_PL_01_01_R1_GG0 | Poland | Ramets | 68,19 | 79,22 | 0,007 |
| PN_PL_01_04_R1_GG0 | Poland | Ramets | 42,56 | 79,70 | 0,005 |
| PN_PL_01_06_R1_GG0 | Poland | Ramets | 55,69 | 79,56 | 0,005 |
| PN_PL_01_08_R1_GG0 | Poland | Ramets | 51,31 | 78,52 | 0,005 |
| PN_PL_01_09_R1_GG0 | Poland | Ramets | 53,38 | 79,36 | 0,005 |
| PN_PL_01_11_R1_GG0 | Poland | Ramets | 35,94 | 80,50 | 0,006 |
| PN_PL_01_12_R1_GG0 | Poland | Ramets | 52,06 | 79,72 | 0,005 |
| PN_PL_01_13_R1_GG0 | Poland | Ramets | 48,31 | 79,94 | 0,005 |
| PN_PL_01_15_R1_GG0 | Poland | Ramets | 49,00 | 79,65 | 0,005 |
| PN_PL_01_18_R1_GG0 | Poland | Ramets | 55,75 | 79,64 | 0,005 |
| PN_PL_01_19_R1_GG0 | Poland | Ramets | 47,63 | 80,75 | 0,006 |
| PN_PL_01_20_R1_GG0 | Poland | Ramets | 40,69 | 79,45 | 0,005 |
| PN_PL_01_21_R1_GG0 | Poland | Ramets | 51,50 | 79,58 | 0,005 |
| PN_PL_01_26_R1_GG0 | Poland | Ramets | 34,88 | 80,27 | 0,006 |
| PN_CZ_01_13_P0_WW0 | C.Republic | Ortets | 36,94 | 78,98 | 0,006 |
| PN_CZ_01_25_P0_WW0 | C.Republic | Ortets | 44,88 | 79,10 | 0,006 |
| PN_CZ_01_33_P0_WW0 | C.Republic | Ortets | 49,06 | 79,36 | 0,005 |
| PN_CZ_01_39_P0_WW0 | C.Republic | Ortets | 46,13 | 78,50 | 0,005 |
| PN_CZ_01_52_P0_WW0 | C.Republic | Ortets | 52,50 | 79,17 | 0,005 |

| | | | | | |
|---|---|---|---|---|---|
| PN_DE_01_10_P0_WW0 | Germany1 | Ortets | 67,25 | 61,88 | 0,006 |
| PN_DE_01_24_P0_WW0 | Germany1 | Ortets | 62,38 | 80,19 | 0,005 |
| PN_DE_01_35_P0_WW0 | Germany1 | Ortets | 64,31 | 79,25 | 0,005 |
| PN_DE_01_38_P0_WW0 | Germany1 | Ortets | 57,69 | 80,02 | 0,007 |
| PN_DE_01_44_P0_WW0 | Germany1 | Ortets | 57,25 | 79,09 | 0,005 |
| PN_ES_01_05_P0_WW0 | Spain | Ortets | 49,69 | 78,39 | 0,004 |
| PN_ES_01_15_P0_WW0 | Spain | Ortets | 52,13 | 77,69 | 0,005 |
| PN_ES_01_16_P0_WW0 | Spain | Ortets | 43,94 | 78,73 | 0,005 |
| PN_ES_01_22_P0_WW0 | Spain | Ortets | 52,19 | 78,69 | 0,005 |
| PN_ES_01_23_P0_WW0 | Spain | Ortets | 53,44 | 79,46 | 0,005 |
| PN_FR_01_01_P0_WW0 | France1 | Ortets | 51,69 | 79,81 | 0,005 |
| PN_FR_01_14_P0_WW0 | France1 | Ortets | 61,56 | 79,54 | 0,005 |
| PN_FR_01_20_P0_WW0 | France1 | Ortets | 85,50 | 79,82 | 0,006 |
| PN_FR_01_22_P0_WW0 | France1 | Ortets | 48,56 | 80,41 | 0,005 |
| PN_FR_01_30_P0_WW0 | France1 | Ortets | 52,88 | 79,53 | 0,005 |
| PN_FR_02_20_P0_WW0 | France2 | Ortets | 50,38 | 79,56 | 0,005 |
| PN_FR_02_26_P0_WW0 | France2 | Ortets | 56,19 | 80,31 | 0,005 |
| PN_FR_02_35_P0_WW0 | France2 | Ortets | 53,63 | 80,51 | 0,005 |
| PN_FR_02_39_P0_WW0 | France2 | Ortets | 44,44 | 80,35 | 0,005 |
| PN_FR_02_43_P0_WW0 | France2 | Ortets | 48,06 | 79,22 | 0,004 |
| PN_IT_02_07_P0_WW0 | Italy2 | Ortets | 65,13 | 78,92 | 0,005 |
| PN_IT_02_09_P0_WW0 | Italy2 | Ortets | 53,81 | 79,04 | 0,006 |
| PN_IT_02_17_P0_WW0 | Italy2 | Ortets | 51,81 | 79,74 | 0,006 |
| PN_IT_02_40_P0_WW0 | Italy2 | Ortets | 57,19 | 79,79 | 0,006 |
| PN_IT_02_44_P0_WW0 | Italy2 | Ortets | 49,44 | 79,13 | 0,006 |
| PN_NO_01_14_P0_WW0 | Norway | Ortets | 55,56 | 79,63 | 0,005 |
| PN_NO_01_18_P0_WW0 | Norway | Ortets | 52,50 | 79,14 | 0,005 |

| | | | | | |
|---|---|---|---|---|---|
| PN_NO_01_19_P0_WW0 | Norway | Ortets | 57,63 | 79,58 | 0,005 |
| PN_NO_01_21_P0_WW0 | Norway | Ortets | 65,13 | 79,54 | 0,005 |
| PN_NO_01_23_P0_WW0 | Norway | Ortets | 64,50 | 79,11 | 0,005 |
